# End-to-end Learning of Safe Stimulation Parameters for Cortical Neuroprosthetic Vision

**DOI:** 10.1101/2025.01.23.634543

**Authors:** Burcu Küçükoğlu, Bodo Rueckauer, Jaap de Ruyter van Steveninck, Maureen van der Grinten, Yağmur Güçlütürk, Pieter R. Roelfsema, Umut Güçlü, Marcel van Gerven

## Abstract

Direct electrical stimulation of the brain via cortical visual neuroprostheses is a promising approach to restore basic sight for the visually impaired by inducing a percept of localized light called ‘phosphenes’. Apart from the challenge of condensing complex sensory information into meaningful stimulation patterns at low temporal and spatial resolution, providing safe stimulation levels to the brain is crucial. We propose an end-to-end framework to learn optimal stimulation parameters (amplitude, pulse width and frequency) within safe biological constraints. The learned stimulation parameters are passed to a biologically plausible phosphene simulator which takes into account the size, brightness, and temporal dynamics of perceived phosphenes. Our experiments on naturalistic navigation videos demonstrate that constraining stimulation parameters to safe levels not only maintains task performance in image reconstruction from phosphenes but consistently results in more meaningful phosphene vision, while providing insights into the optimal range of stimulation parameters. Our study presents a stimulus-generating encoder that learns stimulation parameters (1) satisfying safety constraints, and (2) maximizing the combined objective of image reconstruction and phosphene interpretability with a highly realistic phosphene simulator accounting for temporal dynamics of stimulation. End-to-end learning of stimulation parameters this way enables enforcement of critical biological safety constraints as well as technical limits of the hardware at hand.

## 1 Introduction

Globally, approximately 40 million people suffer from blindness, and 250 million are visually impaired (Ackland et al., 2017). Neuroprostheses that bypass the damaged visual pathway and stimulate the visual cortex could aid them by inducing a rudimentary form of ‘phosphene’ vision. Phosphenes are point-like flashes of light evoked by electrical stimulation (Brindley and Lewin, 1968), occurring in the visual field with respect to electrode locations based on the retinotopic organisation of the visual cortex. Concurrent stimulation of multiple electrodes can elicit a pattern of multiple phosphenes forming basic shapes (Chen et al., 2020) that may, when configured properly, enable a patient to improve their navigation ability and interaction with the environment (Fernandez, 2018). The prosthesis controller has multiple degrees of freedom available, including the location, duration, amplitude, frequency, and pulse width of stimulation (Foroushani et al., 2018; Niketeghad and Pouratian, 2018). Designing an encoder to determine these stimulation patterns based on input from visual surroundings can be phrased as a constrained optimization problem: the chosen stimulation parameters should optimize functional vision (measured by the performance on a task like navigation) while staying within physiologically safe limits. A safe, effective and sustainable neuroprosthesis should activate neural tissue without damage to the tissue itself or the electrodes providing the stimulation (Merrill et al., 2005). Determining these stimulation parameters can be challenging because their effect on perception is complex and not yet fully understood. For instance, repeated stimulation at a single electrode can lead to habituation effects (Schmidt et al., 1996), and simultaneous stimulation of multiple electrodes can combine in nonlinear ways (Fernandez et al., 2021; Bosking et al., 2018). The present work addresses the challenge of determining stimulation parameters with two major contributions. First, we make use of a novel realistic phosphene simulator for cortical prostheses to incorporate the effect of various stimulation parameters and phosphene properties into the encoder optimization. Second, we constrain the learned stimulation parameters to biologically safe values while maintaining meaningful perception.

While previous research has focused on direct mechanistic image translation from camera footage to stimulation patterns (Wang et al., 2021), often using a single kind of feature, such as edges, segmentation maps, or keypoint detection (Dagnelie et al., 2007; Parikh et al., 2013; Vergnieux et al., 2014), others treated this compression challenge as an end-to-end optimization problem (de Ruyter van Steveninck et al., 2022; Küçükoğlu et al., 2022; Granley et al., 2022a) to learn encoders that produce the desired stimulation patterns or parameters. However, de Ruyter van Steveninck et al. (2022); Granley et al. (2022a) excluded temporal aspects of the stimulation by focusing on single images, and de Ruyter van Steveninck et al. (2022); Küçükoğlu et al. (2022) focused less on biological aspects. None of these studies included safety limits regarding neural stimulation.

Recently, van der Grinten et al. (2024) developed a phosphene simulator modeling the stimulation’s effect on perception over time based on a range of psychophysical and neurophysiological findings, and provided an initial demonstration of this differentiable simulator’s capability for end-to-end learning of stimulation parameters on videos of moving digits and with safety-constrained ranges on static images. Our work extends this study in three ways. First, instead of focusing on amplitude only, we explicitly investigate also the effect of stimulation parameters like pulse width and frequency that influence stimulation duration and therefore determine the total amount of charge deposited in the tissue. Second, we apply safety-constrained end-to-end stimulus optimization by constraining the maximal input charge via individual constraints on the values of investigated stimulation parameters. Finally, the optimization application is extended to a highly relevant context, namely to an urban navigation setting with real-life video sequences.

Our end-to-end pipeline consists of a deep neural network encoding stimulation parameters of amplitude, pulse width and frequency of stimulation for a video sequence. The biologically plausible phosphene simulator (van der Grinten et al., 2024) transforms these stimulation patterns into phosphenes by mapping them to their locations in the visual field while modeling their characteristics like size, brightness and temporal dynamics (brightness decay and stimulus habituation). A decoder network uses this simulated phosphene vision (SPV) to reconstruct the original video sequence. Optimization of the whole pipeline occurs via the minimization of an objective composed of the reconstruction error (difference between original and reconstructed images) and the phosphene regularization error (difference between original and phosphene images). Therefore, the optimization is steered towards the reconstruction of the entire video sequence via end-to-end learning of relevant features, rather than an upfront specification of relevant features.

The contribution of our work consists in the inclusion of biological safety constraints, the focus on temporal aspects of stimulation, the simultaneous optimization along multiple dimensions in stimulus space, and the use of real-life video data, all with the aim of facilitating the use of phosphene generation models in real-world settings. We aim to answer the following questions: Which stimulus parameters (amplitude, pulse width, frequency) contribute most to improved task performance based on objectives of image reconstruction and phosphene interpretability? What value ranges for these parameters are learned as optimal as a result of the end-to-end training pipeline? Can stimulus generation be improved by incorporating detailed phosphene characteristics (dynamics, cortical magnification, habituation) into the optimization process? What effect does a safety constraint have on functional phosphene vision?

### Related work

One of the main challenges of neural stimulation is to come up with an effective and safe stimulation protocol that will enable control of the desired reaction in the neural tissue. When it comes to effectiveness in the context of neuroprosthetics, it is crucial to compress complex sensory information in visual surroundings to a lower resolution in a way that fits the retinotopic organization of the cortex. Evoking the desired percepts with the stimulation induced requires two aspects. First, generating sparse electrical stimulation patterns conveying crucial visual information in a scene. Second, mapping the consequences of electrical stimulation to perceptual effects in the brain.

#### Stimulus encoding

For the first aspect, many initial studies took a direct mechanistic scene processing approach. To highlight the most relevant parts in a scene some relied on cues of saliency (Parikh et al., 2013, 2010; Han et al., 2021), depth (Perez-Yus et al., 2017; McCarthy et al., 2014; Han et al., 2021), segmentation (Sanchez-Garcia et al., 2020; Han et al., 2021), environmental structure (Vergnieux et al., 2014; Sanchez-Garcia et al., 2020), or edge filtering (Vergnieux et al., 2017; Boyle et al., 2001; Guo et al., 2018). These approaches often involved processing based on a single kind of feature, e.g. edges, segmentation maps, with each being shown to be well suited for a separate particular scenario, such as navigation, object detection, or emotion recognition (Lozano et al., 2020). To maximize the benefit of computer vision algorithms, de Ruyter van Steveninck et al. (2022); Granley et al. (2022a) have turned to end-to-end optimization of the stimulation patterns via deep learning, trying to go beyond the need for adaptation to specific context while addressing the compression issue. However, these studies have incorporated neither the encoding of the temporal features in the visual data, nor the temporal dynamics of the phosphene percept that arise from physical characteristics of electrical neurostimulation. The former of these temporal aspects in generating optimal stimulation patterns have been indirectly included in other studies by having a sequential setting via the integration of a reinforcement learning (RL) agent in learning a behavioral task with adaptive phosphene vision (Küçükoğlu et al., 2022) or via using RL in identifying the task-relevant salient features in a scene (White et al., 2022, 2019). Still, none of these studies took the mentioned temporal dynamics due to electrical neurostimulation into account. Involving these temporal dynamics relates to the level of biological modeling in simulating the phosphene vision, as it concerns neural and physical effects of stimulation over time on brain and perception. To be able to incorporate this aspect necessitates going beyond single images as input data. Therefore, to generate optimal stimulus encodings, dynamic processing of the visual stimuli sequentially by integration of information in the temporal domain is crucial, which implies the need for biological realism in modeling the temporal dynamics for simulated phosphene vision.

#### Phosphene simulation

All of the studies above can be improved in terms of taking biological realism into account, which corresponds to the second aspect of evoking desired percepts via induced stimulation. This also closely relates to the topic of safe stimulation, as only biologically-realistic mechanisms can guide safety limitations. To incorporate biological mechanisms in cortical visual prostheses, Granley et al. (2022b) utilized brain-inspired convolutional neural networks to predict percepts emerging as a result of stimulation without the modeling of temporal aspects, while van der Grinten et al. (2024) took a modeling approach including temporal dynamics based on neurophysiological findings by fitting clinical data to estimate phosphene percepts based on stimulation parameters. More accurate modeling of the expected percept enables more effective stimulation protocols to be learned with the end-to-end optimization process. van der Grinten et al. (2024) demonstrated the feasibility of such a simulation on videos of moving digits, and additionally on static naturalistic scenes with a constrained safety range for the stimulation amplitude learned.

#### Safe stimulation

Finding an effective condensation of visual information via accurate modeling of perceptual effects of stimulation by itself is barely sufficient when it comes to effective electrical stimulation. Ensuring levels of safety is of utmost importance, which, so far, has not been the priority of computer vision or end-to-end optimization studies focusing on processing of naturalistic sensory information for visual neuroprostheses. With respect to neural stimulation as a general topic, many studies focused on finding safe stimulation protocols that avoid tissue damage while enabling sustainable use of electrodes (Merrill et al., 2005). Shannon (1992) recognized charge density and charge per phase as factors influencing tissue damage which allows for distinguishing between stimulation levels that are tissue damaging or not. Cogan et al. (2016) identified the influence of other factors like pulse frequency, duty cycle and current density on tissue damage. In terms of overpotential of an electrode, the extra potential required to drive electrochemical reactions in tissue beyond its equilibrium potential that may damage the tissue or electrode if too high, Merrill et al. (2005) mentioned the influence of charge per pulse, waveform type, stimulation frequency, electrode’s material, geometric area and roughness.

For visual neuroprostheses, inducing phosphenes comes with a trade-off that necessitates balancing the electrical charge delivered in-between stimulation efficacy and safety goals. On the one hand, a sufficiently high charge per second is needed to exceed the threshold of minimum current required to invoke a phosphene perception. On the other hand, the charge per pulse should be sufficiently low for the safety of electrodes in order to prevent an intolerable overpotential (Foroushani et al., 2018; Merrill et al., 2005). The perception threshold depends on parameters like amplitude, pulse width and frequency of stimulation, as well as train length, inter-train interval, pulse waveform and polarity, electrode locations, impedance, depth, size, and their distance in between, number of stimulation electrodes and repetition of stimulation in certain directions (Foroushani et al., 2018). Pio-Lopez et al. (2021) reports most commonly used satisfactory values used in the literature for these parameters. Among these, we can mention for stimulation amplitudes a range of 1–100 *μ*A as necessary to induce phosphenes (Fernández et al., 2020), for frequency a common use of 200 Hz, and for pulse duration 200 *μ*s or 400 *μ*s in the case of intracortical electrodes (Pio-Lopez et al., 2021). Cohen (2007) reports also a 200 *μ*s pulse width for intracortical electrodes, based on (Schmidt et al., 1996). The same goes for (Rajan et al., 2015), where also a frequency of 300 Hz was used and an amplitude range of 10–100 *μ*A tested for chronic intracortical microstimulation, which was demonstrated to be safe by (McCreery et al., 2010). Kim et al. (2016) analyzed effects of stimulating with pulse width of range 50–400 *μ*s, frequency of range 50-1000 Hz and amplitude between 0–100 *μ*A. McCreery et al. (2002) experimented with prolonged stimulation with ranges of 20-160 *μ*A for amplitude, 50-150 *μ*s per phase pulse width for a biphasic phase and 50 Hz frequency. Chen et al. (2014) investigated effects of chronic intracortical microstimulation in ranges of stimulation amplitude 10–100 *μ*A with 300 Hz frequency and 200 *μ*s phases of a biphasic pulse. Fernandez et al. (2021), whose clinical data was used to fit the model of the biological simulator we used, investigated effects of charge per phase on probability of phosphene perception for frequency values of 100-300 Hz and pulse width of 100-800 *μ*s. They demonstrated lower charge per phase required for 300 Hz compared to 200 or 100 Hz, as well as for 170 *μ*s compared to 100 *μ*s, then 400 *μ*s and finally 800 *μ*s relatively. They also tested stimulus amplitudes in the range of 1-128 *μ*A or used a fixed amplitude of 90 *μ*A in their experiments. Based on these findings, default values to provide to the biologic simulator for stimulation parameters of frequency, pulse width and amplitude were taken as 300 Hz, 200 *μ*s and 100 *μ*A respectively.

These are commonly used in the literature as reasonable, safe and effective values. However, there are many more configurations of stimulation parameters that can be provided, and whose effect has not been tested yet in terms of their efficacy. Particularly, there is still a missing link in finding both effective and safe robust stimulation protocols for neuroprosthetic vision, in the context of naturalistic temporal scenes while taking care of the temporal dynamics of the visual data involved. This study aims to address this gap, by searching for optimal stimulation within a reasonable range of stimulation parameters that would satisfy safety constraint requirements through learning a stimulus generator automatically, which saves time in the case of systems that need to be calibrated for thousands of electrodes. Such an approach provides an advantage over the current clinical practice where individual perception thresholds are mapped first to select slightly above stimulation values for each electrode in a time-consuming and repetitive manual procedure. Our modeling and optimization based approach takes advantage of a range of biological and physical aspects and their temporal interactions instead to simultaneously determine stimulation parameters of all electrodes. Additionally, the combined modeling of the stimulation parameters of amplitude, pulse width and frequency provides higher control over the phosphene appearance. Finally, this study goes beyond existing computational approaches with its contribution in taking safety into consideration.

## 2 Methods

### 2.1 End-to-end deep learning pipeline

Our end-to-end training pipeline for learning optimal safe stimulation parameters for a cortical neuroprosthesis is composed of three architectural components: an encoder, a phosphene simulator and a decoder (Fig. 1). The encoder is a trainable deep neural network that transforms a video sequence into three stimulation vectors for each frame, each vector with a size equal to the number of phosphenes. The stimulation vectors represent the stimulation parameters of interest, namely amplitude, pulse width and frequency. The encoder can be constrained to output safe stimulation levels for these parameters. Next, a phosphene simulator transforms these stimulation parameters into plausible visual percepts of phosphenes. Finally, a trainable phosphene decoder network takes as input this visual phosphene percept to learn to reconstruct the original input sequence to inform the training pipeline about the quality of the phosphene percept. The whole pipeline is trained end-to- end to optimize the parameters of the encoder and decoder networks by minimizing a combined loss term composed of a reconstruction loss measuring the difference between the original sequence and its decoded reconstruction, and a phosphene regularization loss measuring the difference between the original sequence and its visual percept outputted by the phosphene simulator. The goal is to end up with a trained encoder that provides an optimal stimulus generation mechanism, given the constraints of the biological system and the hardware used, and accounting for the temporal dynamics of continuous stimulation.^1^

**Figure 1:**
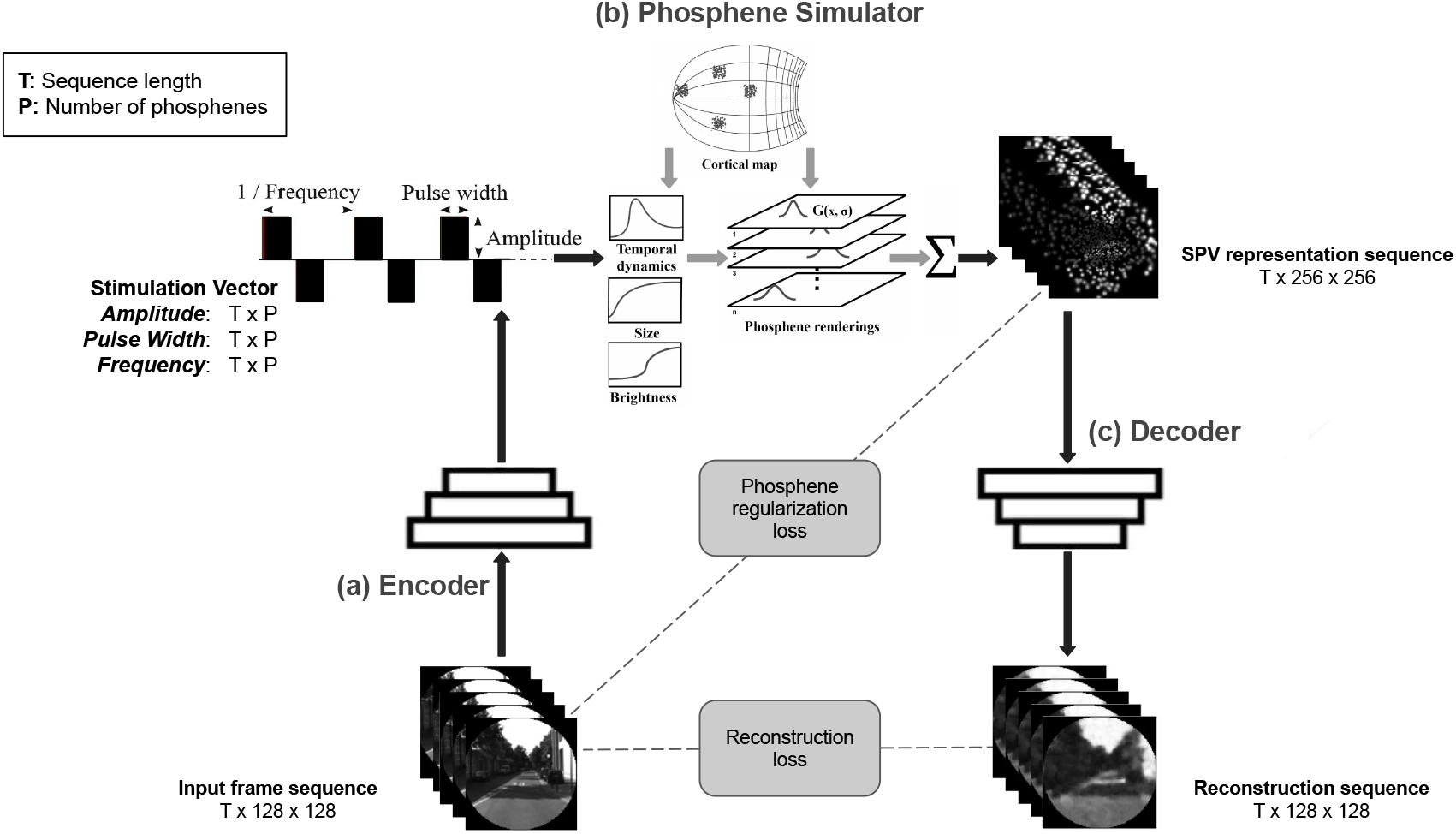
Pipeline for end-to-end learning of biologically plausible safe stimulation parameters for neuroprosthetic vision and its components. (a) An encoder converts video input into stimulation vectors per frame, containing stimulation parameters needed to induce phosphenes via the electrical stimulation of the cortex. (b) A phosphene simulator captures the perceptual effect of the stimulation provided, by modeling phosphene characteristics like temporal dynamics, size and brightness and mapping phosphenes to their given locations in the visual field via Gaussian activation maps G(x,*σ*) to sequentially create a simulated phosphene vision (SPV) representation of size 256 × 256 for each frame in the sequence one by one. (c) A decoder attempts to reconstruct the original input sequence from the SPV representation sequence. The output of the decoder can be used to compute a reconstruction loss, which can be combined with the phosphene regularization loss to drive the whole pipeline towards safe stimulation parameters for optimally generated phosphenes.

### 2.2 Stimulation encoder

The encoder takes as input a video sequence of *T* grayscale frames of size 128 × 128. The encoder produces up to three stimulation vectors per frame, one for each stimulation parameter: stimulation amplitude *I*_stim_ [A], pulse width *t*_p_ [s], and frequency *f* [Hz]. Each encoder output is of size sequence length × number of phosphenes, providing a stimulation parameter value for each phosphene in each frame. As for the network architecture, we compared two choices commonly used with temporal data, as described in the following paragraphs.

#### 3D convolutional encoder

The first architecture which was used is given by a 3D convolutional network that allows for simultaneous spatio-temporal processing. The 3D convolutional encoder (Fig. 2) includes four convolutional blocks, each block including a 3D convolution layer, followed by a 3D BatchNorm and a leaky ReLU activation. Output from the sequential application of these four convolutional blocks are flattened for the last two dimensions of the data. It is then passed through a separate linear layer for each of the stimulation parameters outputted by the encoder, with output size matching the number of electrodes. A final activation function is applied after each linear layer, along with a scaling factor depending on whether the output is constrained for safety or not. This network takes input data of shape *N* × *C* × *T* × *H* × *W*, where *N* is the batch size, *C* the channel size, *T* the time dimension (i.e., the sequence length), *H* height and *W* width of the frame, with kernels sliding over the 3 dimensions of size *T, H* and *W*. The architecture is inspired by the encoder for video representation learning in (Zhao et al., 2017), and builds on the one used in (van der Grinten et al., 2024) for dynamic encoding of moving digits. We have made slight modifications, such as the removal of max pooling layers, whose impact was compensated with larger strides in the preceding convolutional layers, to allow for more learning in the model rather than relying on fixed operations (Springenberg et al., 2015). The model consists of 3.2M parameters in the case of three stimulation vectors generated per frame and 1.1M parameters in the case of one stimulation vector generated per frame.

**Figure 2:**
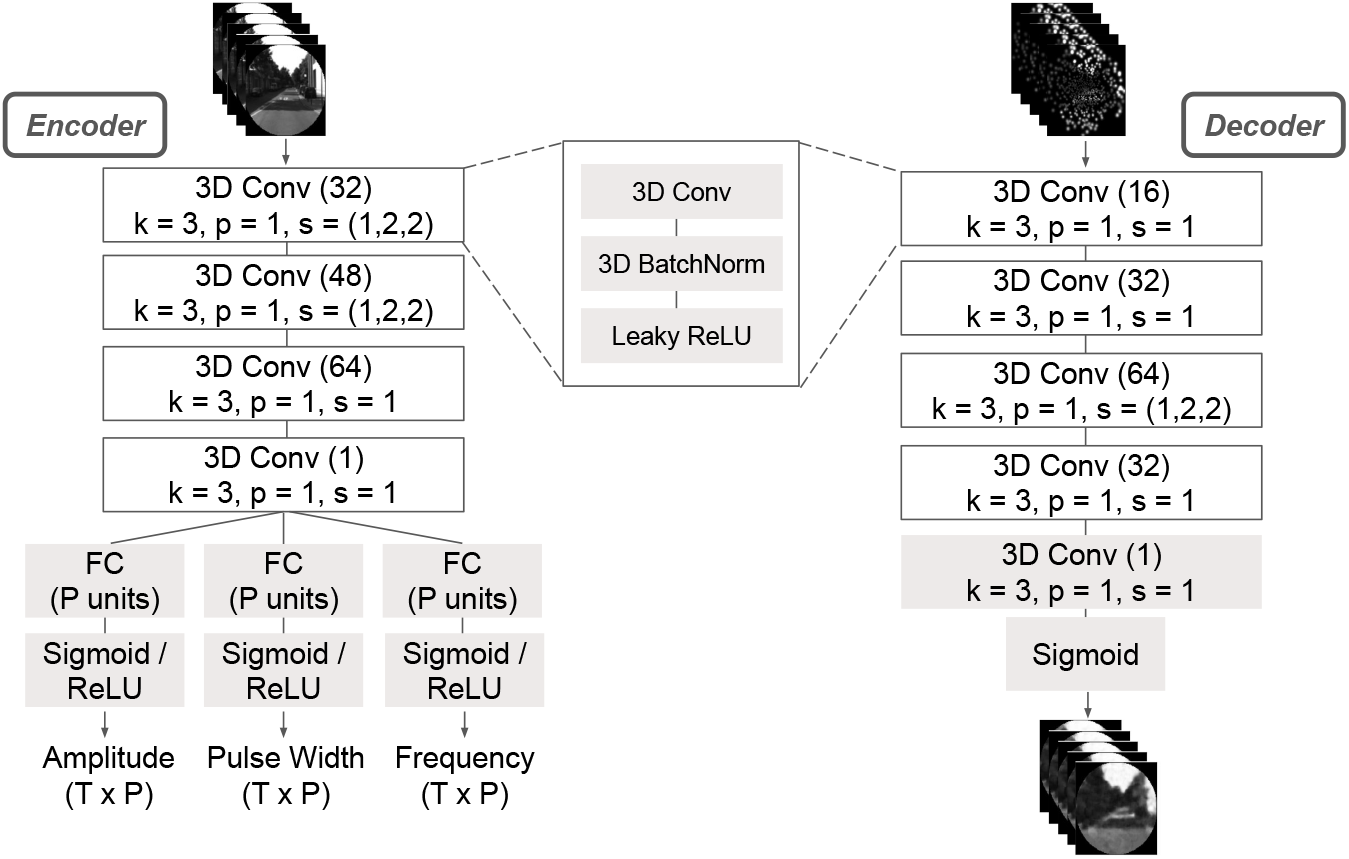
Architectural details of the 3D convolutional encoder (left) and the matching 3D convolutional decoder (right). Grey boxes represent individual layers, white boxes are layer blocks. Kernel size, padding and stride for convolutions are indicated by *k, p* and *s*, respectively. Within parentheses, the number of output channels for convolutions or output sizes for fully connected (FC) layers are provided. The final activation function for the encoder is a sigmoid if constraints on stimulation parameters are applied, or a ReLU if not. A bias is used only in the final convolutional layer of the decoder.

#### Recurrent encoder

The recurrent encoder (Fig. 3) includes three convolutional blocks, followed by two residual blocks and another convolutional block. Each of the convolutional blocks include a 2D convolution layer, followed by a 2D batch normalization and a leaky ReLU activation. Each of the residual blocks include a 2D convolution layer, a 2D batch normalization, a leaky ReLU, another 2D convolution layer and 2D batch normalization sequentially. The final convolutional block has an output channel matching the sequence length. Output from the sequential application of these blocks are flattened for the last two dimensions of the data, and is then passed through a linear layer, and then an LSTM layer. The LSTM output is passed through a separate linear layer for each of the stimulation parameters generated by the encoder, with output size matching the number of electrodes. A final activation function is applied after each linear layer, along with a scaling factor depending on whether the output is constrained for safety or not. This network takes input data of shape *N* × *T* × *H* × *W* with channel dimension treated as the time dimension due to the use of single-channel grayscale images. The architecture is a combination of a modified version of the 2D encoder in (van der Grinten et al., 2024) with slight modifications like removal of some residual blocks and of max pooling layers, whose effect was included with changes in the strides of convolutional layers, and of an LSTM layer, with linear layers before and after. The model has a total of 9.6M parameters in the case of three stimulation vectors generated per frame and 7.6M parameters in the case of one stimulation vector generated per frame. The hidden state is initialized to zero at the beginning of each video sequence batch.

**Figure 3:**
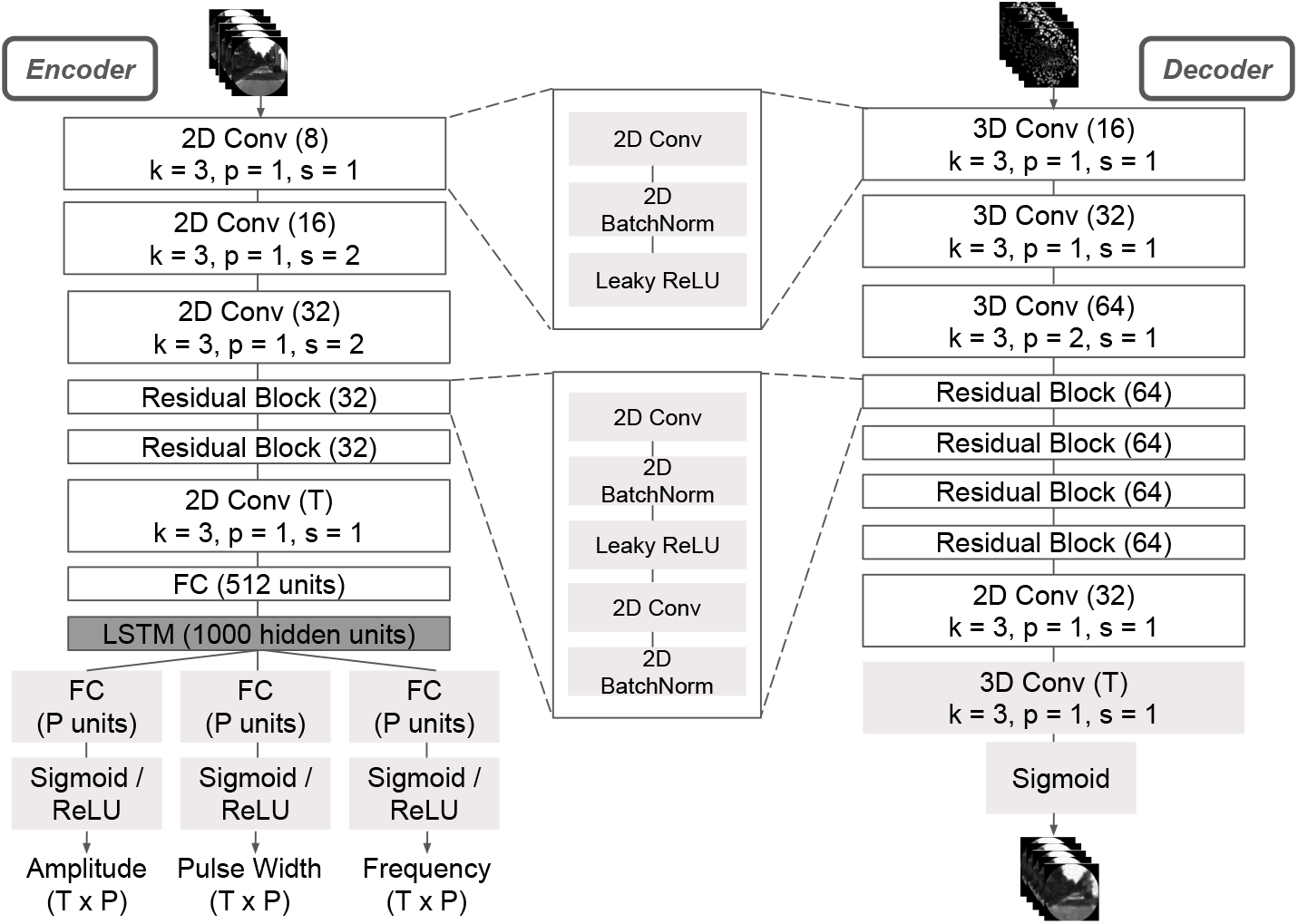
Architectural details of the recurrent encoder (left) and the matching 2D convolutional decoder (right). Grey boxes represent individual layers, white boxes are layer blocks. Kernel size, padding and stride for convolutions are indicated by *k, p* and *s*, respectively. Within parentheses, the number of output channels for convolutions or output sizes for fully-connected (FC) layers are provided. The final activation function for the encoder is a sigmoid if constraints on stimulation parameters are applied, or a ReLU if not. Biases are used only in the convolutional layers of the residual blocks and the final convolutional layer of the decoder.

### 2.3 Phosphene simulator

The phosphene simulator, which was directly taken from (van der Grinten et al., 2024), captures the perceptual effect of cortical stimulation via the use of differentiable functions. The simulator has two modes of initialization, from which we used the more basic one, where a custom list of phosphene locations within the visual field is provided by the user in terms of polar coordinates, to ensure that the initialization of the simulator is the same across our experiments. The simulator takes as input the stimulation parameters amplitude, pulse width and frequency for each phosphene in a frame, one frame at a time. This requires a vector of size (number of phosphenes) from each of these input parameters. These parameters can be provided as output from the encoder, or can also be given fixed default values across the phosphenes in a frame and across frames. The simulator takes all three stimulation parameters to calculate their combined effect as effective current (charge per second) by the multiplication of pulse width *t*_p_, frequency *f* and the effective component of the amplitude, which is defined as stimulation amplitude *I*_stim_ minus its ineffective component *I*_0_ and a memory trace *Q*:

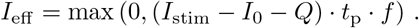

The ineffective component *I*_0_ represents the minimal current for infinite stimulation duration (see Table 1 for values of parameters mentioned throughout this section). The memory trace is a dy-namic update of the stimulation history at each frame, which takes care of the temporal dynamics accounting for habituation effects of brightness due to prolonged or repeated stimulation via *Q*_*t*_ = *Q*_*t*−Δ*t*_ + Δ*Q*_*t*_, where

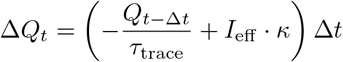

with *τ*_trace_ and *κ* the trace decay and increase rates per second.

**Table 1:** Parameters used in the phosphene simulator.

| Parameter | Description | Value |
| --- | --- | --- |
| $I_0$ | ineffective component of stimulation amplitude | $23.9 \mu\text{A}$ |
| $\tau_{\text{trace}}$ | trace decay rate | $0.995 \text{ s}^{-1}$ |
| $\kappa$ | trace increase rate | $14.0 \text{ s}^{-1}$ |
| $\tau_{\text{act}}$ | activation decay | $1.23 \cdot 10^{-4} \text{ s}^{-1}$ |
| $d$ | relative duration of stimulator to frame | 1 |
| $\lambda$ | slope of sigmoid activation function | $192 \cdot 10^5$ |
| $A_{50}$ | half maximum brightness value | $1.06 \cdot 10^{-7}$ |
| $K$ | excitability constant | $675 \cdot 10^{-6} \text{ A/mm}^2$ |
| $k$ | scaling factor mapping to realistic proportions in cortical distance | 17.3 |
| $a$ | dipole model stability parameter | 0.75 |
| $b$ | dipole model stability parameter | 0.4 |
| $\theta$ | horizontal view angle | $16^\circ$ |

By integrating the effective current over multiple frames, cortical tissue activation *A*_*t*_ is estimated based on a leaky integrator model with updates for each frame given by *A*_*t*_ = *A*_*t*−Δ*t*_ + Δ*A*_*t*_, where

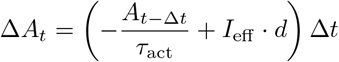

with activation decay per second *τ*_act_ and *d* scaling the duration of stimulation relative to the frames. When activation is greater than the phosphene’s detection threshold, the phosphene is activated with a brightness based on the sigmoidal activation 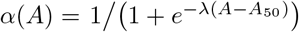, where *λ* is the slope and *A*_50_ signifies the activation value where a phosphene reaches half the maximum brightness. Note that a detection threshold larger than zero is not used in our study despite the simulator allowing for it. This was chosen to ensure same conditions across our experiments as the implementation of a stimulation threshold introduces some degree of variability between phosphenes via sampling of activation thresholds from a normal distribution. Moreover, based on the possibility of subthreshold activation from concurrently stimulated electrodes to cause above-threshold activations in larger areas than predicted by single electrode studies (Bosking et al., 2018), we opted to keep the thresholds at lowest to enable optimization through concurrent stimulation.

Phosphene size *P* is calculated as the ratio *P* = *D/M* between the diameter *D* of activated cortical tissue and the cortical magnification factor *M*. Here, 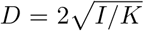 with *I* the stimulation amplitude and *K* the excitability constant, determining how far the spread of activation reaches. The cortical magnification factor *M*, defined in terms of millimeters of cortical surface per degree of visual angle, signifies the relative amount of cortical tissue involved in processing of visual information depending on the eccentricity in the visual field. It is obtained via the dipole mapping model (Polimeni et al., 2006) for phosphene topological mapping, giving the cortical magnification factor

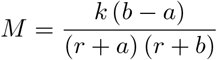

where *r* is eccentricity of the point in the visual field, *k* is a scaling factor for mapping to realistic proportions in cortical distance, and *a* and *b* are parameters controlling singularities of the dipole model. Finally the phosphenes are visualized as Gaussian blobs, where 95% (two standard deviations) of the Gaussian falls within the calculated phosphene size. They are rendered on their initialized locations in the visual field via a meshgrid, after conversion of their coordinates from polar to Cartesian coordinates. Here, phosphene locations outside the horizontal view angle, taken as 16 degrees, are removed. The possibility to apply Gabor filtering in a certain orientation and frequency in the simulator beyond the default Gaussian blobs is ignored in this study for simplicity and to avoid random variation across experiments.

The processing of the simulator at every training step proceeds as follows. At the beginning of each sequence, the simulator is reset. This involves the memory trace, cortical tissue activation and size values to be set to zero just before taking a video input. The phosphene simulator takes as input the stimulation parameters for each frame of the video one by one, therefore being applied as many times as the sequence length at each training step. With every frame, effective charge per second is calculated, then the phosphene states of tissue activation, memory trace, size and brightness are updated via the aforementioned equations based on the current stimulation input and the previous states of the values in time. Later, Gaussian activation maps are generated for phosphenes based on their size and location mapping. Then, phosphene intensities are set via their brightness values wherever the cortical activation is greater than zero (or greater than the detection thresholds if they were to be used), keeping the other phosphenes at zero intensity. At the end, an image with the simulated phosphene representation is returned by mapping intensity values across the Gaussian activation maps. The sequential application of the simulator on each frame in a video in this manner allows for temporal dynamics to be followed over a single video sequence.

### 2.4 Phosphene decoder

The phosphene decoder takes as input a sequence of grayscale SPV representations of size sequence length × 256 × 256. It learns to output a sequence of grayscale reconstructions of the original input sequence of size sequence length × 128 × 128. Depending on the encoder architecture used, the decoder can have two different choices for its architecture. First, a 3D convolutional network for the decoder is used together with a 3D convolutional encoder. Second, a 2D convolutional decoder is used together with a recurrent encoder.

#### 3D convolutional decoder

The 3D convolutional decoder (Fig. 2) includes four convolutional blocks, which include the same components described for the 3D encoder. Output from the sequential application of these four blocks are passed on to another 3D convolutional layer. A final sigmoid activation is applied to get an output between 0 and 1, matching the range of normalized images inputted to the encoder. This architecture is inspired by the decoder from (Zhao et al., 2017), and is taken from the one used in (van der Grinten et al., 2024) for moving digits. The model has a total of 126,001 parameters.

#### 2D convolutional decoder

The 2D convolutional decoder (Fig. 3) includes three convolutional blocks, which include the same components described for the recurrent encoder. After the sequential application of these convolutional blocks, four residual blocks are applied, as described for the recurrent encoder. After the sequential application of residual blocks, another convolutional block is applied. At last a 2D convolutional layer is applied, followed by a sigmoid activation, outputting values between 0 and 1 to match the range of normalized images provided to the encoder. This architecture is taken from (van der Grinten et al., 2024; de Ruyter van Steveninck et al., 2022). The model has a total of 342,538 parameters.

### 2.5 Safety constraints

Current amplitude, pulse width, and pulse frequency determine the total amount of charge deposited in the neural tissue over time:

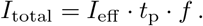

Setting a single limit on the combined term *I*_total_ is not enough because the contribution of an unduly high pulse amplitude *I*_eff_ could be concealed by a small frequency *f*. We therefore set individual constraints on each factor. In the constrained case, a scaled sigmoid function was used to map the encoder’s fully connected layer outputs to the interval [0, *α*], where *α* is the desired safety constraint for the parameter of interest:

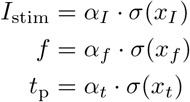

where *x* stands for the output of the fully connected layer for the respective parameter. To prevent pulses from overlapping at high frequencies, the pulse width was specifically limited by setting *α*_*t*_ = 1*/*(2*f*) when learning all stimulation parameters. In the unconstrained baseline case, the stimulus parameters can take on any positive value (implemented by a ReLU activation function in the final layer of the encoder).

## 3 Experiments

### 3.1 Experimental design

In our experiments, we compare the performance of unconstrained against constrained training of all three stimulation parameters (amplitude, pulse width, frequency). Next, only a single stimulation parameter was learned under its constraint while the others were kept fixed at their default values to determine their individual contribution (e.g. amplitude constrained). The constraints were set to 500 *μ*A, 500 *μ*s and 500 Hz to include the most commonly tested values in the literature for amplitude, pulse width and frequency respectively, whereas the fixed default values were chosen as 100 *μ*A, 200 *μ*s and 300 Hz based on commonly used values (please see the Related Work section for an overview). All experiments were conducted both with the 3D convolutional and the recurrent encoder. The reported results are averages over five random seeds.

### 3.2 Dataset

The KITTI Vision Benchmark Suite (Geiger et al., 2012) was used as the dataset, which includes navigation videos in real cities captured at 10 Hz frame rate, hence requiring the frames per second to be set to 10 also in the simulator. Although the dataset included videos of various lengths, we split them into videos of equal length by taking a sequence length of 10 frames, giving one-second-long videos, which also makes the train duration of stimulation 1 s for electrodes active during the entire video. To make use of more data, videos were split into 10 frames by sliding over the long video frames with a step size of 2, causing some overlap in the frames across the 10-frame videos we used. This way, a total of 9339 videos with sequence length 10 was used. With a train-test data split ratio of 9:1, the training dataset is composed of 8405 videos while the test set is composed of 934 videos. Basic preprocesing was applied on the frames, converting them to single-channel grayscale images, center cropping and resizing to 128 × 128 pixels from their varying original sizes, as well as normalizing its values to a range from 0 to 1. Finally a circular mask was applied on square frames to place black pixels everywhere in the corners apart from the central disc in the square, in order to only capture the circular region of the image that would be covered by phosphenes via our phosphene simulator. This ensures that corner regions of the original images are ignored during loss calculation.

### 3.3 Optimization procedure

During training, the encoder and the decoder are trained simultaneously in an end-to-end fashion to minimize a combined loss term composed of a reconstruction loss and phosphene regularization loss (Fig. 1). The reconstruction loss term 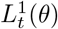 is given by the mean squared error between the original video sequence and the decoded reconstruction sequence, informing us about the phosphene quality via the decodability of the SPV representations. The phosphene regularization term 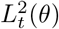 is given by the mean squared error between the sequence of SPV representations and the original input sequence, scaled up to the 256 × 256 phosphene resolution via trilinear interpolation. This loss term informs about the interpretability of the phosphenes. Combining these loss terms yields the objective

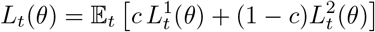

that is to be minimized with 𝔼_*t*_[·] the empirical mean over a set of training samples. The coefficient *c* trades off the reconstruction loss against the phophene regularization loss, which was taken as 0.5 in this study. For training, the Adam optimizer was used with a learning rate of 10^−4^ and a mini-batch size of 2. The networks were trained until the validation loss did not improve further during 15 validation sessions. Validation sessions occurred at every 20 mini-batches of data, while training statistics were saved at every 60 mini-batches of data. Convergence happened usually in a range of 6 to 16 epochs for the constrained case, while it could happen as early as the first epoch for the unconstrained case.

## 4 Results

### 4.1 Comparison of stimulation protocols

We first compare unconstrained and constrained stimulation protocols for the different architectures (Fig. 4). Analysis of statistical t-tests for the loss means shows that the best total loss minimization is achieved by architectures trained without constraints. While the convolutional architecture reaches a similar total loss level to the unconstrained case in the all parameters constrained setting (*p >* 0.05, *N* = 10), the recurrent architecture shows an increased loss (*p* < 0.05, *N* = 10). Constraining all parameters gives better total loss minimization (*p* < 0.05, *N* = 10) than constraining any stimulation parameter alone in both architectures. Similar lowest levels of total loss for both unconstrained and all constrained settings show that we can still reach optimal loss levels despite imposing multiple constraints, showing the advantage of the all constrained case as opposed to one. The all constrained setting gives better loss optimization (*p* < 0.05, *N* = 10) compared to the amplitude constrained setting especially, regardless of loss term or the encoder architecture type analyzed. This specifically demonstrates the benefit of end-to-end framework’s simultaneous optimization of all three parameters as opposed to amplitude only. Constraining amplitude only gives better total loss minimization (*p* < 0.05, *N* = 10) than constraining pulse width or frequency alone in both architectures. These observations suggest that of the three parameters considered here, learning the amplitude is both necessary and sufficient for optimal performance under constraints. If aiming for the two settings that involve amplitude constraining, use of the CNN has advantages (*p* < 0.05, *N* = 10) over the RNN. Pulse width or frequency only constrained settings do not seem to differ in their optimization performance across loss terms and architecture types, and in terms of total loss these two have the worst minimization performance. Yet, in terms of total and reconstruction losses, the RNN seems to be better (*p* < 0.05, *N* = 10) than the CNN in optimizing for the pulse width only constrained setting, while the pulse width constrained RNN is better minimized (*p* < 0.05, *N* = 10) than the frequency constrained CNN. Overall, the CNN encoder architecture seems to make a more positive difference out of all possible cases when desiring a benefit from the move from an unconstrained to a constrained setting. This is especially demonstrated by CNN’s advantages in all constrained and amplitude only constrained cases.

**Figure 4:**
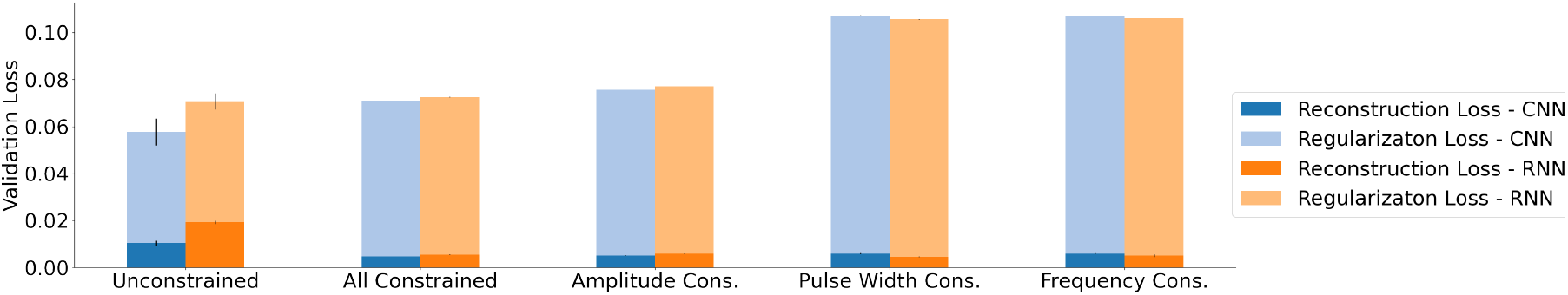
Final values of the means of total validation losses across runs, based on equally contributing components of reconstruction and phosphene regularization losses for different constraint levels and encoder architectures. Error bars show standard error of the mean over runs. Full bars show total loss.

### 4.2 Contribution of loss terms

When we consider the contribution of each loss term and how they compare across experiments, we see that reconstruction loss is actually better minimized (*p* < 0.05, *N* = 10) in the constrained cases. This suggests an advantage of constraints in reconstruction loss optimization. As reconstruction loss term informs us about the phosphene quality via the similarity of reconstructions to the original frames, we could say that constraining the parameters provides better phosphene quality that allows decodability to the original frames. This is understandable, since constraints allow for a reasonable range of stimulation parameters that can only give rise to biologically plausible stimulation effects. Hence, the evoked percepts are also more recognizable. The better minimization of reconstruction loss in the constrained cases show that the reason total loss is lower for the unconstrained case is due to the minimal contribution of the regularization loss.

Phosphene regularization loss is lower (*p* < 0.05, *N* = 10) for the unconstrained case as opposed to each of the constrained cases, signaling a disadvantage of contraints in minimizing regularization loss. The regularization loss term informs about the interpretability of phosphenes via their similarity to the original frames, calculated by pixelwise mean squared error between original frames and SPV representations. Fully minimizing this term is harder because of its calculation based on pixel-wise error and phosphenes having certain locations, so that only in pixels where a phosphene is activated this loss can be minimized. Therefore, covering all phosphene locations with activations would provide the best capability to the network to minimize this loss term. As the network outputs unreasonably high values for amplitude in the unconstrained case, the phosphene sizes are calculated to be larger, hence causing huge Gaussian blobs, sometimes even overlapping with each other. This causes most of the phosphene locations and even the gaps beyond to be lighted up in pixels, making it closer to the original images, hence allowing a better minimization of this loss term. On the other hand, more targeted and sparse activation of phosphenes in the constrained cases does not allow this loss term to be further minimized especially for the black pixels in the SPV representation. While the difficulty of minimizing the regularization loss might lead to questions regarding the role of this term in the optimization, Fig. 5 shows that when optimization lacks the regularization loss term, phosphenes do not seem as sparse or informative for the constrained case, whereas uninformative phosphenes emerge for the unconstrained case as a result of overstimulating parameters, albeit with reconstruction similar to original image. On the other hand, the optimization with only the regularization loss term (Fig. 6) shows the contribution of this term in making phosphenes look informative especially in the constrained case, hence proves useful for our optimization purposes.

**Figure 5:**
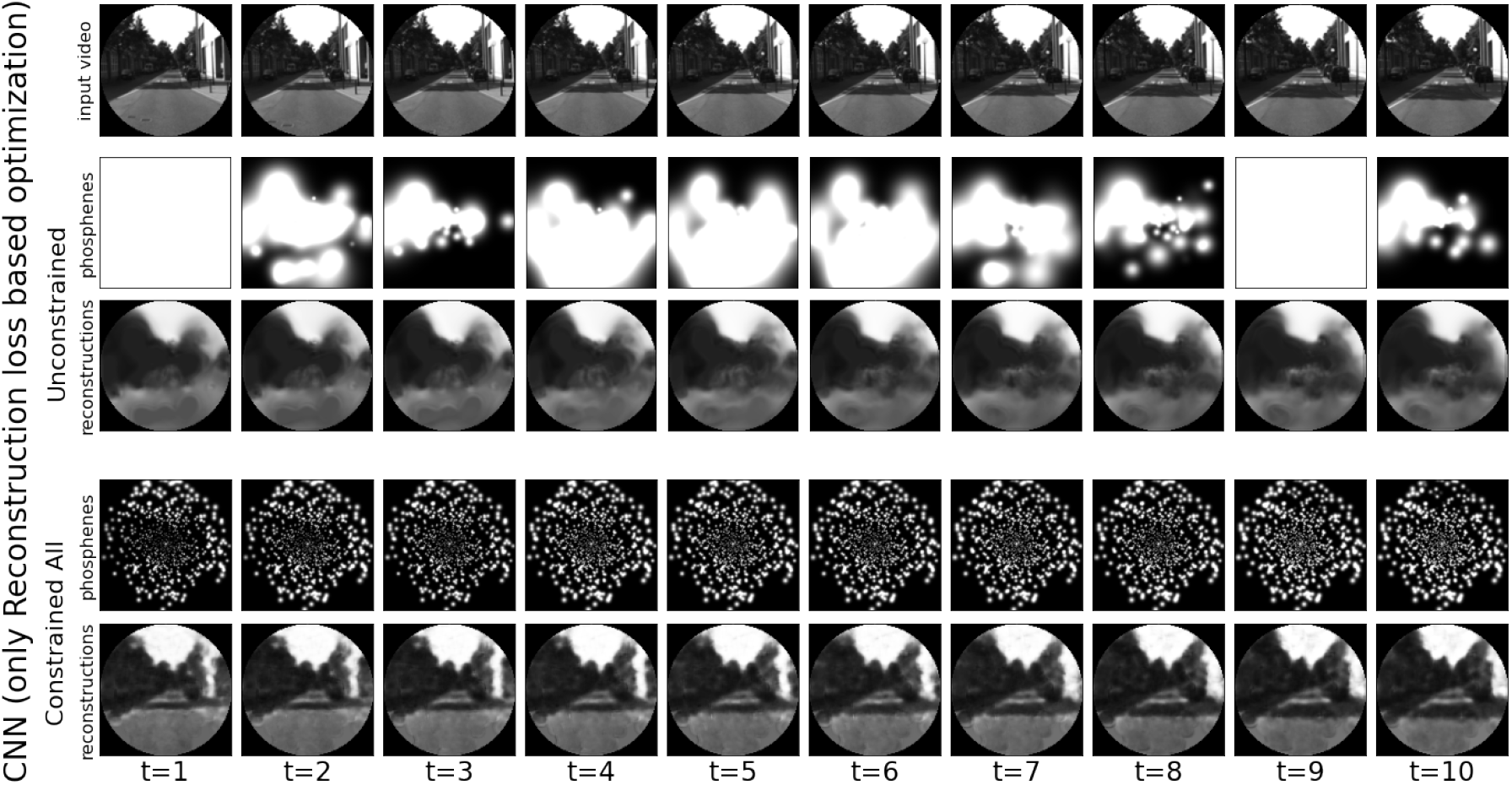
Transformations of original video frames (first row) to phosphenes (second row) and reconstructions (third row) using the CNN architecture in the unconstrained and constrained case with only reconstruction loss in the optimization.

**Figure 6:**
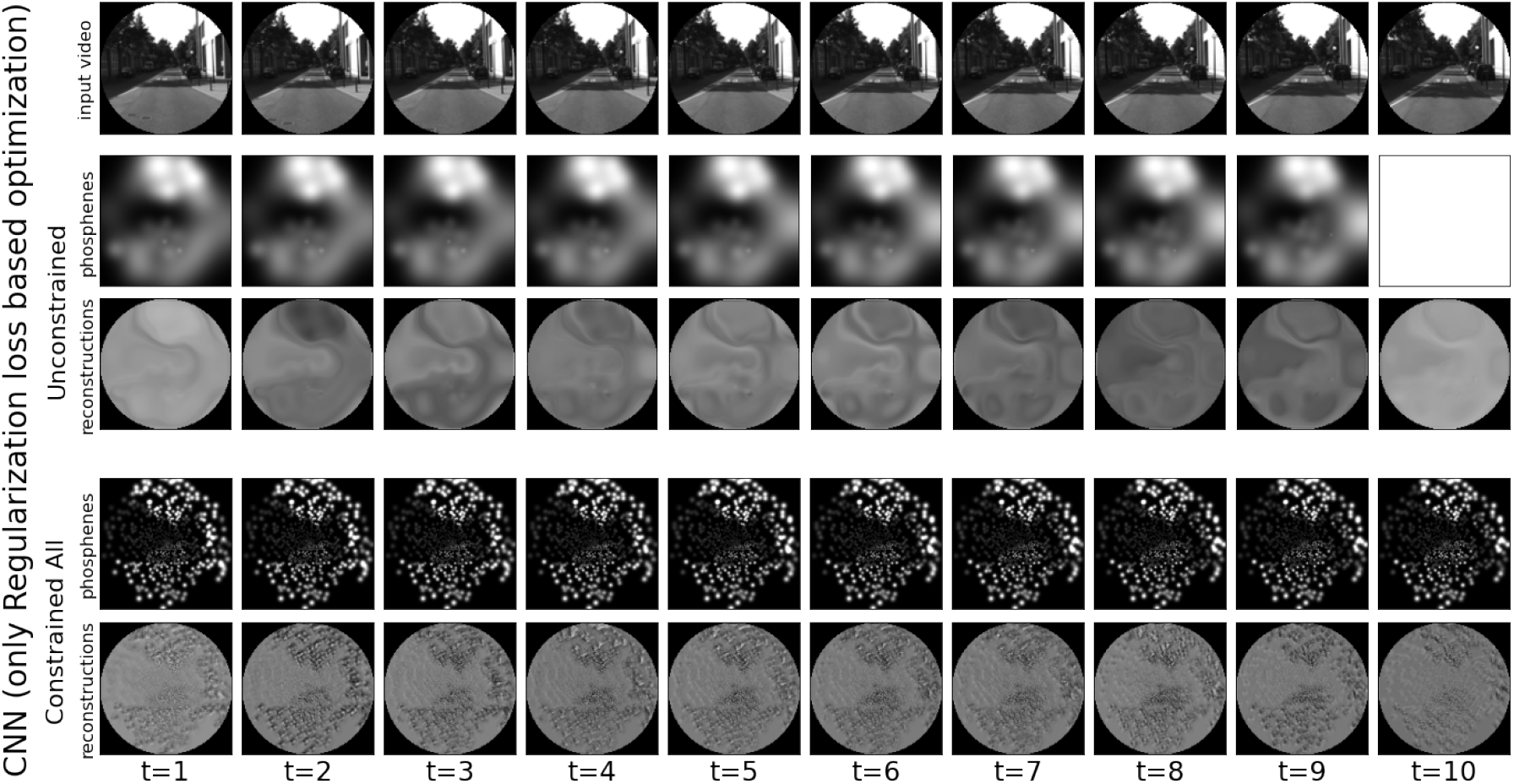
Transformations of original video frames (first row) to phosphenes (second row) and reconstructions (third row) using the CNN architecture in the unconstrained and constrained case with only regularization loss in the optimization.

### 4.3 Inspection of phosphene quality

We next inspect phosphene representations qualitatively. Figures 7 and 8 show the transformation that each frame in the original input sequence goes through via the phosphene vision optimization pipeline, including the SPV representation and the reconstruction of each frame, for the CNN and RNN architectures. The videos of the same transformations are also provided online.^2^ These demonstrations provide a means to inspect the sequential processing of data for each of the experiments conducted. As can be seen, constrained encoders are able to generate informative phosphenes that look similar to the original scenes. The corresponding decoders generate reconstructions that are similar to the original frames. On the other hand, the unconstrained encoders produce blurry and uninformative phosphenes with large blobs.

**Figure 7:**
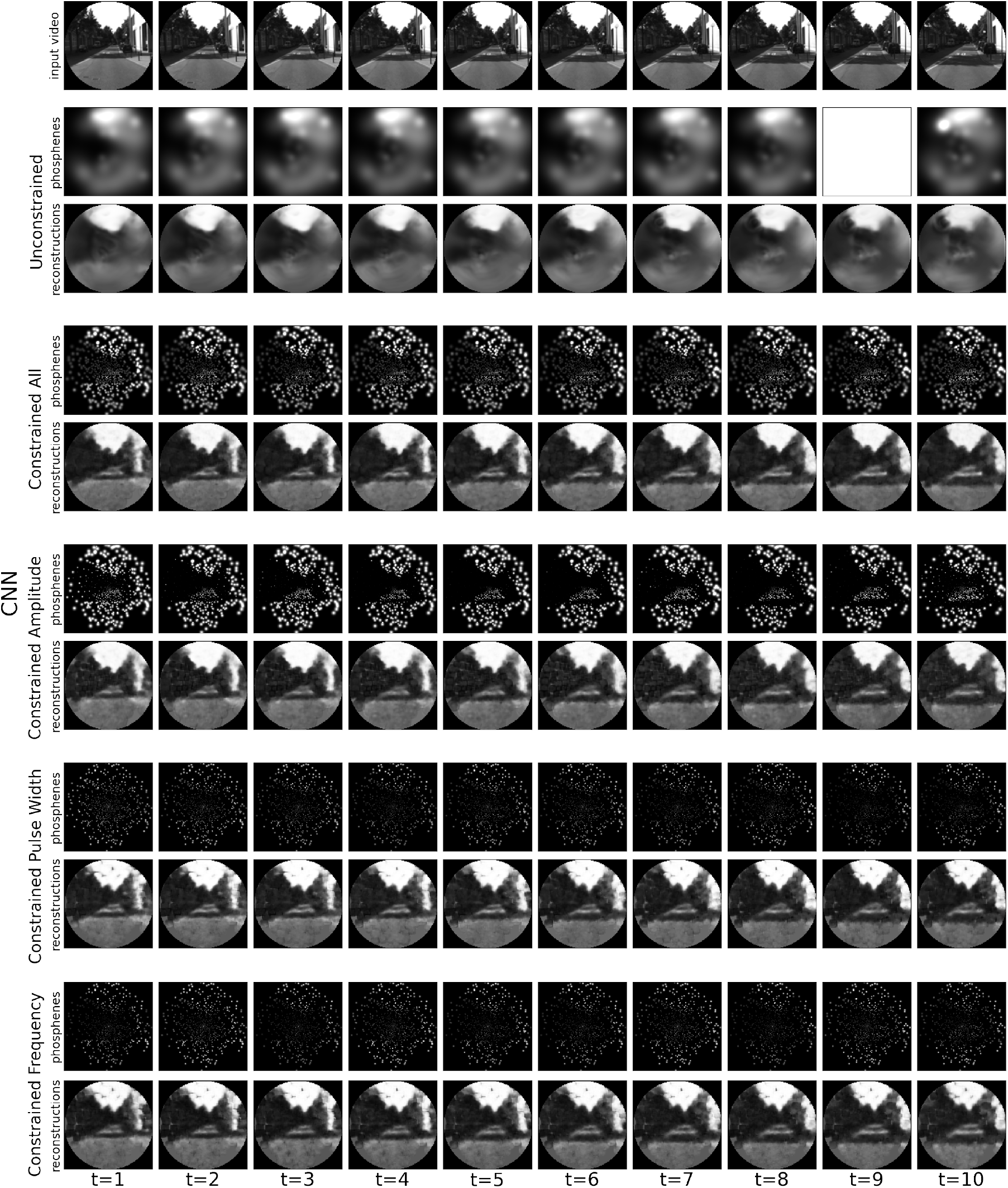
Transformations of original video frames (first row in each subfigure) to phosphenes (second rows) and reconstructions (third rows) with different constraint levels for the CNN architecture.

**Figure 8:**
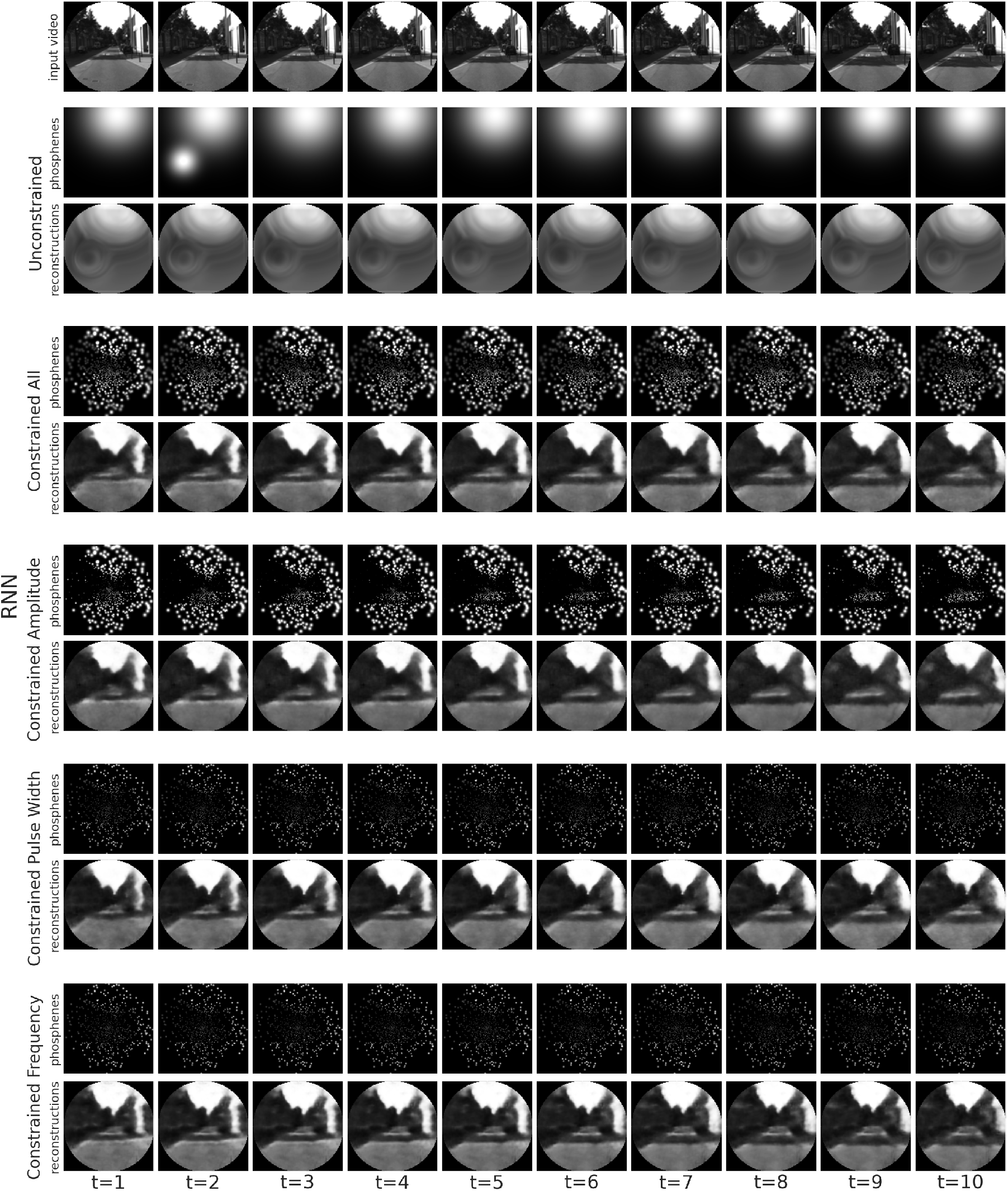
Transformations of original video frames (first row in each subfigure) to phosphenes (second rows) and reconstructions (third rows) with different constraint levels for the RNN architecture.

The different constrained cases produce similarly structured phosphenes, and equally good reconstructions, regardless of the architecture used. The only difference comes with the intensity of phosphenes activated in only pulse width or frequency constrained cases, which are much lower than the cases of only amplitude constrained or all constrained. This is understandable, given the much greater range the amplitude was allowed to be learned, compared to its fixed default value in the cases it wasn’t directly constrained along with other parameters. This is similar for other parameters, as well. However, the gap is the greatest for the amplitude, therefore when amplitude is fixed, the intensity is more likely to be lower.

When we compare all constrained to amplitude constrained, we also see that there is a slight difference in the sense that the all constrained case is more descriptive yet less sparse than the amplitude constrained case. That is, there are more activated phosphenes in the all constrained case, yet more black areas in the amplitude only case. Which case is more desirable depends on the kind of scenes processed, or could also be determined based on biological or hardware requirements that need to be considered e.g. in terms of the total charge per second that can be handled.

### 4.4 Distributions of learned stimulation parameters

To understand the implications of the learned stimulation parameters for biological requirements and hardware capabilities, as well as to inspect how well they align with the findings in the literature, Fig. 9 inspects the distributions of different stimulation parameters generated by the encoder, namely stimulation amplitude, pulse width and frequency, as well as their product, which amounts to the total charge per second (current). We observe a large difference in the learned parameter ranges. The amplitude values learned in the unconstrained case reach tens of amperes, which is unreasonably high for biological systems that were shown to safely tolerate on the order of a few hundred *μ*A for intracortical microstimulation (Rajan et al., 2015; McCreery et al., 1990; Graziano et al., 2002; McCreery et al., 2010). Similarly, pulse width values learned in tens of seconds in the unconstrained case are not feasible for dynamic scenes especially for the 1 s long videos used in the study, with typical values ranging in hundreds of microseconds (Foroushani et al., 2018). Frequency on the other hand goes as low as microhertz in the unconstrained case, which suggests effectively no stimulation, whereas commonly hundreds of Hz are most effective (Foroushani et al., 2018). These observations demonstrate the value of learning the stimulation parameters within safety constraints.

**Figure 9:**
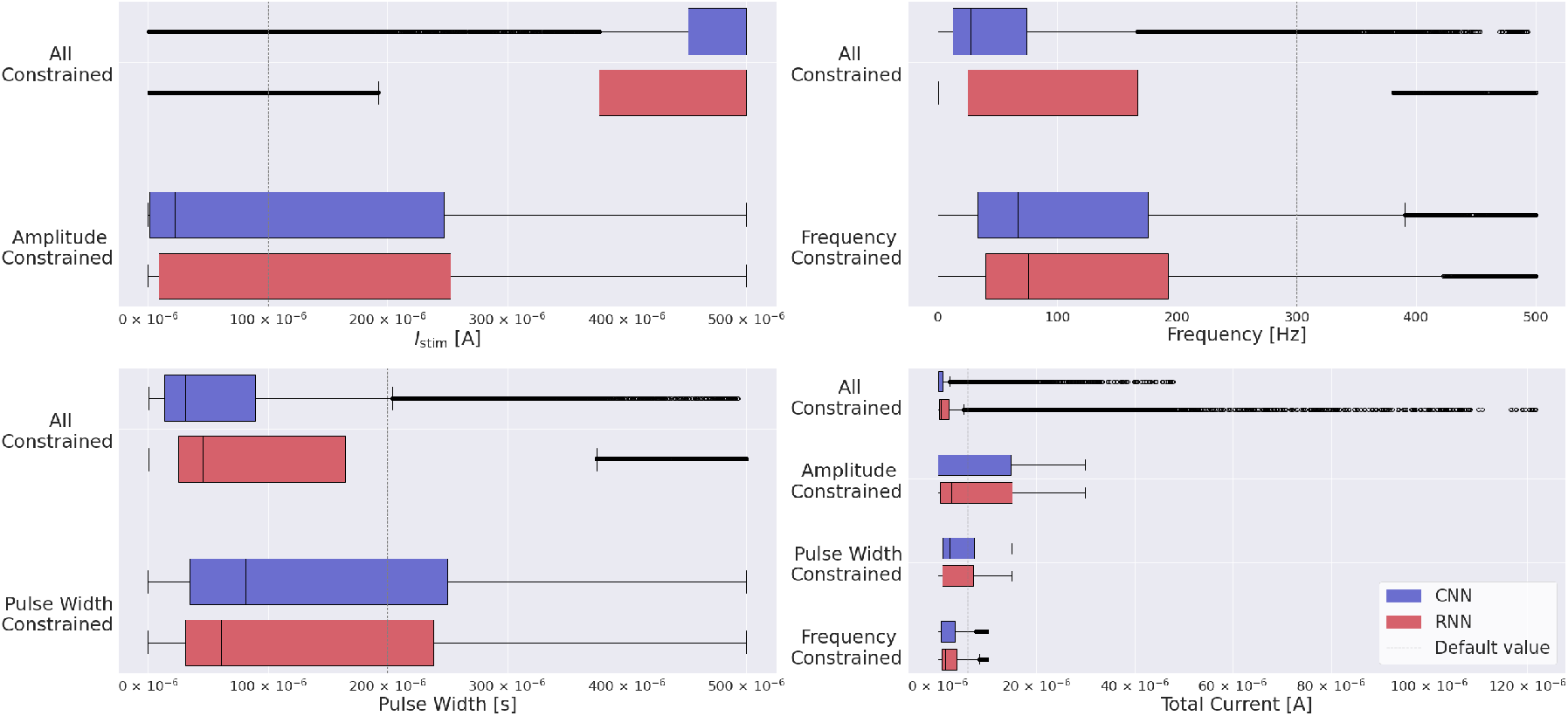
Distribution of stimulation parameters, as well as total current (their simple multiplication corresponding to charge per second) learned in the constrained parameter settings. In each subfigure, the units shown correspond to the unit of values generated by the phosphene simulator for the stimulation parameter of interest. Dashed line represents the default value chosen for each parameter when it is non-trainable, or for current, their multiplication.

The constrained encoders show a tendency for conservative parameter values at the lower end of the allowed range. For pulse width, the majority of values falls below 200 *μ*s with the median being below 100 *μ*s. For frequency, the majority of values falls below 200 Hz with the median being below 100 Hz. This behavior is in line with the relative inefficiency of longer pulse widths and higher stimulation frequencies (Fernandez et al., 2021; Niketeghad et al., 2019; Winawer and Parvizi, 2016) and highlights the importance of using a realistic phosphene simulator for learning the appropriate values for efficient stimulation.

An exception is given by the amplitude, where the learned values cluster around the upper end of the spectrum in the case where all parameters are constrained. In contrast, for the amplitude constrained case, the relative gathering of amplitude values on the lower end of the range shows how the learned values may depend on the default values of the fixed stimulation parameters in the single parameter constrained cases. Similarly, in the all constrained case, pulse width and frequency values adjust to high amplitude values learned, by gathering relatively on the lower end of the range compared to the single parameter constrained cases. The analysis becomes clearer as we see the distributions of current for the constrained experiments. Despite the high amplitude values leading to a greater range of current in the all constrained setting, in the majority of the cases, the resulting total current is actually lower than when only one parameter is learned. Hence, constraining all parameters can be beneficial, especially since it pushes the encoder to minimize their multiplication, which corresponds to their combined effect to the simulator. The generally higher values in the single parameter constrained cases must be caused by the default values, because the encoder minimizes the value of the remaining free parameter. Together, these observations suggest that it may be possible to maintain the same level of perception with lower overall control energy by using learned dynamic stimulation parameters rather than static default values.

On the other hand, due to the extreme parameter values generated in the unconstrained case, the current values in this setting go beyond the order of milliamperes, which can cause seizures upon simultaneous stimulation of multiple electrodes (Fernandez et al., 2021), while current on the order of microamperes suffices to evoke phosphenes with penetrating electrodes (Beauchamp et al., 2020; Bak et al., 1990). These make the current values learned without constraints highly infeasible and thus demonstrate the value of learning safe stimulation parameters in a constrained setting.

## 5 Discussion

In this work we demonstrate the use of an end-to-end pipeline to learn safe stimulation parameters for optimized neuroprosthetic vision, given physiological safety constraints. For this purpose, an encoder is trained on video data to output optimal electrical stimulation parameters of amplitude, pulse width and frequency, whose effect on neural tissue and visual perception are modelled via a differentiable phosphene simulator. This simulator also accounts for temporal dynamics while modeling perceptual phosphene characteristics like size and brightness. Our experiments demonstrate the benefit of constraining the encoder output for learning safe stimulation parameters while retaining optimality in phosphene quality, as otherwise ineffective and unsafe stimulation parameters and biologically implausible phosphene patterns are learned. By incorporating biological safety constraints, a focus on temporal aspects and the use of naturalistic video data, this work contributes to the field by showcasing the capability of end-to-end deep learning to determine optimal safe stimulation parameters for effective realistic phosphene vision.

Whereas the unconstrained end-to-end learning of stimulation parameters leads to the best performance in terms of optimizing for the machine learning objective, the results show that they provide low quality of phosphene vision due to overstimulation, hence biologically implausible phosphenes. Surprisingly, learning stimulation parameters through a CNN while imposing safety constraints on all three stimulation parameters can reach similar loss minimization performance, with the additional benefit of providing the most descriptive phosphenes. Constraining all three stimulation parameters simultaneously leads to better loss performance and informativeness of phosphenes than when constrained singularly. Still, a constraint on the amplitude alone provides more value for optimization performance and informative phosphenes than a singular constraint on parameters of pulse width or frequency. These findings suggest amplitude as the central parameter to constrain for biologically plausible and safe end-to-end learning of phosphene vision, while emphasizing advantages of simultaneous constraints on multiple stimulation parameters for highest benefits.

An important aspect of optimizing for neuroprosthethic vision is handling temporal data while taking into account habituation effects of continuous stimulation. Given the modeling of temporal dynamics in the biophysically realistic simulator, learning stimulation parameters end-to-end is a promising way to compensate for neural habituation. We compared 3D CNNs and recurrent encoder architectures for dealing with temporal data and found an advantage of CNNs for the all constrained and amplitude constrained cases, whereas the recurrent encoder only brought an advantage in the pulse width constrained case. Advantage of CNNs could be due to the short window size adapted for the KITTI sequences, since for training purposes we used a fixed video length, which is long enough to accommodate phosphene dynamics, in line with train durations in clinical studies (Chen et al., 2014; Kim et al., 2016; Rajan et al., 2015; Pio-Lopez et al., 2021). We expect an advantage of (gated) RNNs once the model is applied to longer sequences as in the actual neurostimulation context, given their capability to process long term dependencies (Sutskever et al., 2014; Hochreiter and Schmidhuber, 1997). Hence, uninterrupted video input in an online setting might increase the difference between recurrent and 3D CNN architectures.

Despite using a realistic phosphene simulator, some ambiguity remains in the level of biological detail due to lack of consistent findings in the literature with regard to the effect of some factors in safe stimulation, which inevitably influenced some design choices. One such limitation concerns the pulse shape, as the simulator is agnostic to the waveform. Values taken in modeling of the simulator are from findings in the literature related to biphasic pulses, yet details like whether they are anodic or cathodic first are not specified. By only modeling the effective charge per second rather than pulse design, the simulator also neglects details like interphase interval. Accurate inclusion of such details could lead to more effective learning of optimal stimulation patterns. Another possible limitation is the simplified modeling of tissue activation via the leaky integrator, which might lead to an insufficiency to capture some non-linear dynamics. Also, current modeling data depends mostly on single electrode settings as the studies mostly report on that instead of multi electrode settings, which may in fact give rise to unexpected unrecognizable percepts by merging phosphenes into larger sizes (Beauchamp et al., 2020). Additionally, while the current depiction of perceptual experience is modeled based on detection thresholds, actual detection in patients is influenced by attention and training (Fernandez et al., 2021). Finally, the current pipeline does not take into account eye movements, which leads phosphene locations to move in space accordingly unlike the given fixed locations in this study. Incorporation of such further details in future studies could lead to the generation of more effective stimulation parameters.

When it comes to safety limits, many factors beyond the stimulation parameters directly constrained here have been shown to affect tissue damage, such as current, electrode area, duty cycle, current density and electrode size (Cogan et al., 2016). While this work did not include a direct constraint on the current but rather an indirect one through the product of constrained stimulation parameters, keeping track of its value demonstrated the use of learning all three stimulation parameters in minimizing it. Whereas the maximum total current possible in our experiments due to the chosen parameter constraints or default values were 125 *μ*A, 30 *μ*A, 15 *μ*A, 10 *μ*A for the cases of all constrained, only amplitude, pulse width, frequency constrained settings respectively, the majority of the current values obtained as a result of parameter optimization were below 15 *μ*A or even 10 *μ*A. These values are in line with threshold currents reported for phosphene perception via microstimulation with minimum values of 1 *μ*A (Lycke et al., 2023), 1.8 *μ*A (Orlemann et al., 2024), 1.9 *μ*A (McCreery et al., 2010; Schmidt et al., 1996), and majority of values being below 25 *μ*A (Schmidt et al., 1996) or 20 *μ*A (McCreery et al., 2010; Orlemann et al., 2024), with a typical range of 10–20 *μ*A (Cohen, 2007) or a general range of tens of *μ*A with values extending to 100 *μ*A (Fernández et al., 2020; Fernandez et al., 2021). Future work could include a direct constraint on total charge per second provided across all electrodes, rather than focusing on finding optimal values for single electrodes, which would serve sparsity purposes. Other factors like electrode size or area were included indirectly via the model underlying the simulator, which was fit on physiological data (Fernandez et al., 2021; Horton and Hoyt, 1991; Schmidt et al., 1996) with parameters adapted to the type of electrodes. Investigating the effect of varying electrode properties on stimulus optimization could provide a more comprehensive understanding of optimal stimulation settings.

In the current study, separate stimulation parameters were generated for each electrode at each frame in the video sequence. Such frequent adaptation of stimulus parameters may neither be required nor technically feasible. We chose this setup here to demonstrate the optimization capability of an end-to-end learning pipeline in full generality. The effect of including technical constraints of a real prosthesis prototype will be investigated in future work.

Though informed by the literature, the default values for the non-trainable stimulation parameters and the constraints on stimulation parameters set in this study may to some extent seem arbitrary. Yet, the insights provided by this work stem from the presence or absence of constraints rather than their specific value, demonstrating the capacity of end-to-end learning of stimulation parameters based on safety constraints. Future work will focus on the inclusion of realistic constraints especially in multi-electrode settings, for which little data is available as of yet. While future work can provide more accurate learning of safe stimulation parameters by incorporation of improved knowledge on biologic details or hardware constraints such as the spatiotemporal granularity at which stimulation electrodes can be controlled, the current study lays the groundwork for safe optimization of neuroprosthethic vision in naturalistic video settings by taking into account temporal dynamics.

## Acknowledgments

This work has received funding from the European Union’s Horizon 2020 research and innovation programme under grant agreement No 899287 (project NeuraViPeR). This publication is also part of the project Dutch Brain Interface Initiative (DBI2) with project number 024.005.022 of the research programme Gravitation which is (partly) financed by the Dutch Research Council (NWO). This publication is also part of the project INTENSE with project number 17619 of the Crossover programme.

## Footnotes

1 Code for the end-to-end training pipeline is available at https://github.com/burcukoglu/dynaviseon.git. Updated code of the biologically plausible phosphene simulator for the end-to-end training pipeline in this study is available at both https://github.com/burcukoglu/dynasim.git and the original repository for the simulator https://github.com/neuralcodinglab/dynaphos.git.

2 Videos of transformation of original frames through the phosphene vision pipeline via trained networks can be seen in https://www.youtube.com/playlist?list=PLOsc1mo6nsPfP8JPHmHrYzCOMMGz-zDB7.

## Notes

### Competing Interest Statement

The authors have declared no competing interest.

